# ADHD: relation between cognitive characteristics and DAT1 / DRD4 dopamine polymorphisms

**DOI:** 10.1101/452805

**Authors:** Pinto María Cristina, Ávila Jorge Enrique, Polanco Angela María, Vásquez Rafael Antonio, Arboleda Humberto

## Abstract

Attention deficit hyperactivity disorder (ADHD) is a clinical and diagnostic heterogeneous picture. This study analysed the association of functional polymorphisms in DAT1 VNTR 3’ UTR and DRD4 VNTR Exon III candidate genes, and the neuropsychological characterisation of attention and executive functions of a group of children with ADHD vs. controls. 32 patients and 51 controls were selected. The DAT1 10-repeat allele appeared more frequently in the two groups (cases: 0.93/control: 0.82), showing an OR: 2.5 (IC 95%: 0.684-9.133; p: 0.158). In DRD4, the 4-repeat allele shows the highest occurrence (cases: 0.62/controls: 0.60). None of the markers presented a significant association after a direct analysis, but the DRD4 7-repeat marker showed a positive risk when performing a Bayesian logistic analysis (coefficient: −1.69; OR: 5.39 CI 95%: 1.167-40.97). On the other hand, when considering association with cognitive performance, a positive risk for processing speed and attention tasks was identified.

---

Attention deficit hyperactivity disorder (ADHD) is an early onset neurological, cognitive and behavioral disorder with heterogeneous clinical manifestations, which hinders its diagnosis and differentiation from other common conditions in childhood (Banaschewski et al, 2003; Bará, Vicuña, Pineda Henao, 2003; Bradshaw, 2001; Curatolo et al, 2008; Cinnamon, Willcut, DeFries y Pennington 2007; Faraone & Biederman, 1998; Muñoz, Palau, Salvadó y Valls 2006; Pineda, Puerta, Aguirre, García y Kamphaus, 2007).

Heritability of ADHD is ≈ 0.80, shows the important role of genetics in the etiology of the disorder (Faraone & Biederman, 1998; Curatolo et al, 2008; Biederman et al, 1995; Cordell & Clayton, 2005; Laird & Lange, 2006). It is common to find multigenerational families with a clear pattern of genetic transmission, demonstrated even in Colombian population (Arcos-Burgos, Castellanos, Lopera et al, 2002; Palacio, Castellanos, Pineda et al, 2004). Most cases occur sporadically without a clear Mendelian inheritance pattern, being more compatible with a multifactorial model (Faraone, Perlis, Doyle et al, 2005). The high heritability includes several loci of genetic susceptibility, and the gene-environment interactions contributing to ADHD phenotype variability, although the balance between these various factors is still unknown (Banaschewski et al, 2003). By using unbiased whole-genome amplification methodologies, several susceptibility loci have been established for ADHD in the human genome; 4p16, 5p13, 6q14, 6q26, 7p13, 9q33, 11q25, 13q33, 15q15, 16p13, 17p11, 20q13 (Bakker, van der Meulen, Buitelaar et al., 2003; Heiser, Friedel, Dempfle et al., 2004), also including dopamine receptors (DRD4 & DRD5), dopamine transporter (DAT), serotonin receptor 1B and SNAP-25 (Banaschewski et al, 2003; Curatolo et al, 2008). Most of these approaches point to the identification of candidate genes, from molecular perspectives (Faraone et al, 2005), but without offering explanatory models that relate pathway from genotypes to behavioral manifestations (Rommelse et al. 2008).

The typical findings that have shown a certain consistency in ADHD are those related to the role of dopamine, from pharmacological perspectives and their relation to the processes of modulation of neuronal signal-to-noise ratio (Bush, 2010), as an approach to understand their influence on attentional processes as a basis for the cognitive and behavioral manifestations of ADHD).

The dopamine transporter is highlighted, by modulating the availability of the protein in certain areas of the brain (Heinz, Goldman, Jones et al, 200; Van Dyck, Malison, Jacobsen et al, 2000; Stein, Waldman, Sarampote et al, 2005), with the use of knockout mouse models, Giros *et al*. (1996) showed that the alteration of the dopamine transporter gene (DAT1) led to a hyperdopaminergic phenotype that included spontaneous hyperlocomotion (Miller y Madras, 2002). Some of its variants modulate the response to treatment in patients with ADHD (Faraone et al, 2005; Roman et al., 2002; Cheon, Ryu, Kim y Cho; 2005; Roman, Rohde y Hutz, 2004; Polanczyk, et al, 2005; Cordell y Clayton, 2005), particularly, by being the target of psychostimulants methylphenidate, the main therapeutic agent in ADHD (Heiser et al, 2004).

The frequency of the different polymorphic variants changes widely among populations with different ethnic origin (Kang, Palmatier y Kidd, 1999; Mitchell, Howlett, Earl, et al, 2000; Vieyra, Moraga, Henriquez, Aboitiz y Rothhammer, 2003); however, multiple genetic association studies have consistently found that a variant of this polymorphism (the 10-repeat allele of a variable number tandem repeat (VNTR) in the 3 untranslated region (UTR) with 480pb) confers a moderate risk for the development of ADHD (Barr, Xu, Kroft et al, 2001; Chen, Chen, Mill et al, 2003; Cook, Stein, Krasowski et al, 1995; Curran, Mill, Tahir et al, 2001: Gill, Daly, Heron, Hawi y Fitzgerald, 1997; Mill, Ronald et al., 2005; Todd, Huang, Smalley et al, 2005; Purper-Ouakil, et al, 2005). Several recent meta-analysis support this conclusion for both case-controls and intrafamily studies (Faraone et al, 2005; Hedebrand, Dempfle, Saar et al, 2005) becoming one of the polymorphisms with greater consistency in neuropsychiatric diseases (Greenwood, Alexander, Keck et al, 2001, Ogdie, Bakker, Fisher et al, 2005; Bannon, 2005).

On the other hand, the dopamine receptor D4 gene (DRD4), due to its pharmacological characteristics, cerebral expression and role in dopaminergic pathways, has been widely studied for genetic association with several neuropsychiatric, behavioral disorders (Wong, Buckle, Van Tol, 2005; Lopez et al., 2005), as and behavioral characteristics collectively called ‘Novelty Seeking’ (Faraone y Biederman, 1998; Lopez et al., 2005; Currier, Grandy, Gerhardt y Glaser, 2009), which gathers aspects such as impulsivity, exploration, excitability and irritability, observed between those with ADHD (Faraone, Biederman, Weiffenbach et al. 1999). DRD4 receptor with 7-repeat (7R) variant shows an unmitigated response to dopamine, finding high rates of this allele in children with ADHD compared with ethnicity and gender-matched controls (LaHoste *et al.* 1996). These results suggest that the distribution of this receptor in the brain plays an important role in cognitive and emotional functioning. Our analysis aims to explore the possible risk association between the presence of functional polymorphisms in candidate genes (DAT1 VNTR 3 ‘UTR and DRD4 VNTR Exon III) and ADHD, as well as to determine the association between these polymorphisms and cognitive performance in tests of executive function and attention in Colombian children population.

## Methods

### Participants

This study used a case-control design, with children from Bogotá–Colombia. The cases (n = 32) were recruited from Hospital La Misericordia (HOMI) child psychiatric service, with ADHD diagnosis established according to DSM-IV-TR (APA, 2002). Healthy controls (n = 50), from public schools, with no diagnosis in clinical records or diagnostic scales. All subjects age ranging from 6 to 16 years. The protocol was approved by the Universidad Nacional-Medicine Faculty and HOMI ethical committee.

The legal guardians of minors signed informed consent, and participants provided verbal approval (informed assent). None of the participants had comorbidity with learning disorders or other behavioral and neurological conditions affecting development (autism, TB1, epilepsy, neurological or genetic disorders, use of psychoactive substances). Participants who obtained an IQ of <70 on the Wechsler Intelligence Scale for Children WISC IV (Wechsler, 2007) were excluded.

Subjects were asked to provide saliva samples from mouth rinse and participate in cognitive evaluation.

### Cognitive Assessments

The neuropsychological evaluation protocol included standardized measurements of multiple cognitive domains, selected by their ability to evaluate cognitive functions associated with ADHD, mainly related to attentional processes and executive function:

*Wechsler Intelligence Scale — [WISC IV]* (Wechsler, 2007): it allowed identifying the level of general intellectual functioning, and evaluating skills such as focal (selective and sustained) attention, processing speed, and working memory. The necessary subtests for obtaining the scale of total IQ were used, but the test of numbers and letters was replaced by the arithmetic subtest to consider performance in these skills, including this value in global scale, as well as in working memory index.

*Child neuropsychological Assessment – [ENI]* (Matute, Rosselli, Ardila y Ostrosky-Solís, 2007): selected subtests included cancellation of drawings and letters to analyze visual tracking skills and selective attention.

*Stroop Color and Word test* (Golden, 1999): Assessment of resistance to interference-inhibition.

*The Swanson, Nolan and Pelham Questionnaire [SNAP-IV]* (Bussing, Fernandez, Harwood, Hou, Wilson, Eyberg y Swanson, 2008 y Grañana, Richaudeau, Robles Gorriti, O’Flaherty, Scotti, Sixto, et al. 2011) was used to assess clinical symptoms and manifestations, and severity with parents and teachers.

### Genetic analysis

Genotyping of the VNTR 3’UTR, dopamine transporter (DAT1) gen and VNTR Exon III polymorphisms of the dopamine receptor 4 (DRD4) gen was performed, using standardized methods previously described in some international publications by the Neuroscience Group from Universidad Nacional de Colombia (Forero, Arboleda, Yunis, Pardo y Arboleda, 2005; Forero, et al, 2006).

For VNTR, the following primer sequences were used: DRD4: F: 5’GCGACTACGTGGTCTACTCG 3’; R: 5’AGGACCCTCATGGCCTTG 3’. DAT1: F: 5’TGTGGTGTAGGGAACGGCCTGAG 3’; R: 5’CITCCTGGAGGTCACGGCTCAAGG3’. These genetic markers were selected based on the analysis of previous evidence of polymorphisms that represent risk of ADHD and a possible theoretical association with neurocognitive functioning.

### Statistical analysis

Chi-square analysis was carried out for gene frequencies of alleles. For all neuropsychological tests, measures were transformed to z-scores, considering the values of the mean and standard deviation for comparison and participants were divided into groups differentiated by the presence or not of the allele of interest. Comparisons were carried out through the Fisher’s exact test, considering the size of the sample and the expected frequencies. Subsequently, an analysis of odds ratio (OR) and relative risk (RR) was performed, including a confidence interval value of 95% (CI). Finally, a logistic Bayesian and logistic regression analysis model was adjusted based on a priori information for strengthening risk analysis. The threshold for statistical significance was P <0.05. All analyzes were performed using SPSSv19 (IBM, 2010) and R (R Core Team, 2010) statistical programs.

## Results

**Table 1.**
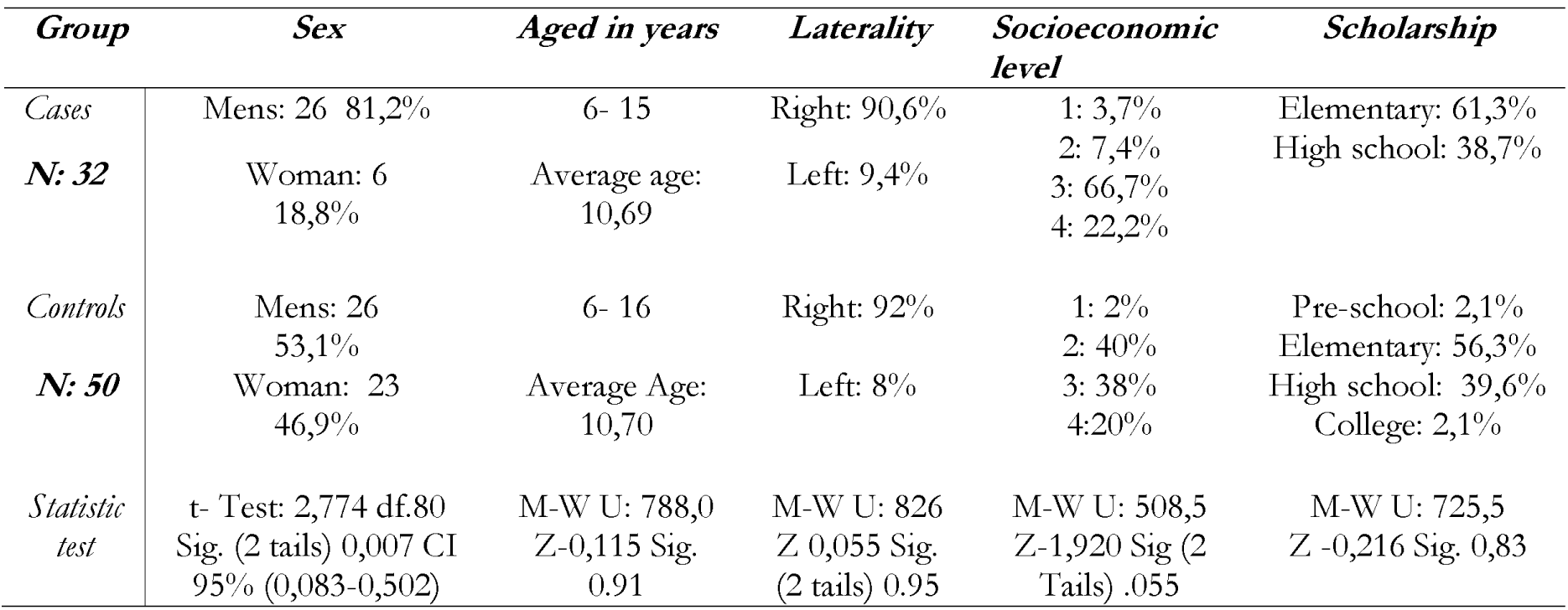
*Characteristics of the analyzed population*

It was found that both groups share demographic characterization, with an average age of 11, are mostly right-handed and belong to socioeconomic level 3. It was also found that the only variables for which finding significant differences is possible between the groups are related to sex, where a different proportion of men and women is seen, which is consistent with the findings of incidence in ADHD (Cardo & Servera-Barceló, 2005; Cornejo et al., 2005; Pineda, Lopera, palacio y Ramírez, 2003; Pineda et al, 1999).

**Table 2.**
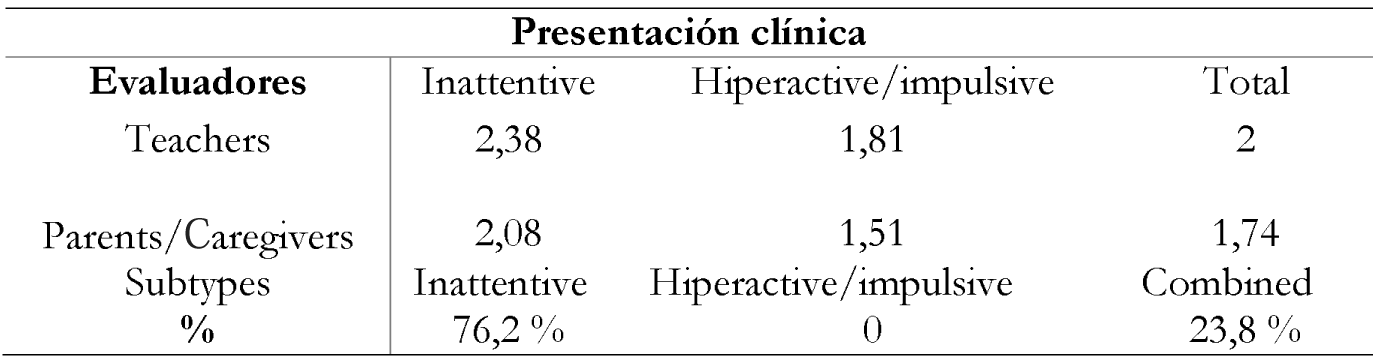
*SNAP IV average scale scores and derivative variants.*

Within the cases group, and based on the analysis of the average of SNAP IV test scales (Bussing et al., 2008; Grañana et al., 2011), participants were grouped into two clinical manifestations: inattentive, being the most frequent with 76.2%, and mixed or combined subtype with 23.8%. No scores that exceed the cutoff point of 95% (1.66) for the inattentive subtype, exclusively, were obtained. It was also observed that parents assigned lower intensity to general symptoms compared with teachers, and, for both evaluators, the manifestations that occurred with greater severity (scores> 2) are associated with inattention. Table 3 shows results related to allele frequencies; populations were in HW equilibrium for both markers.

**Table 3.**
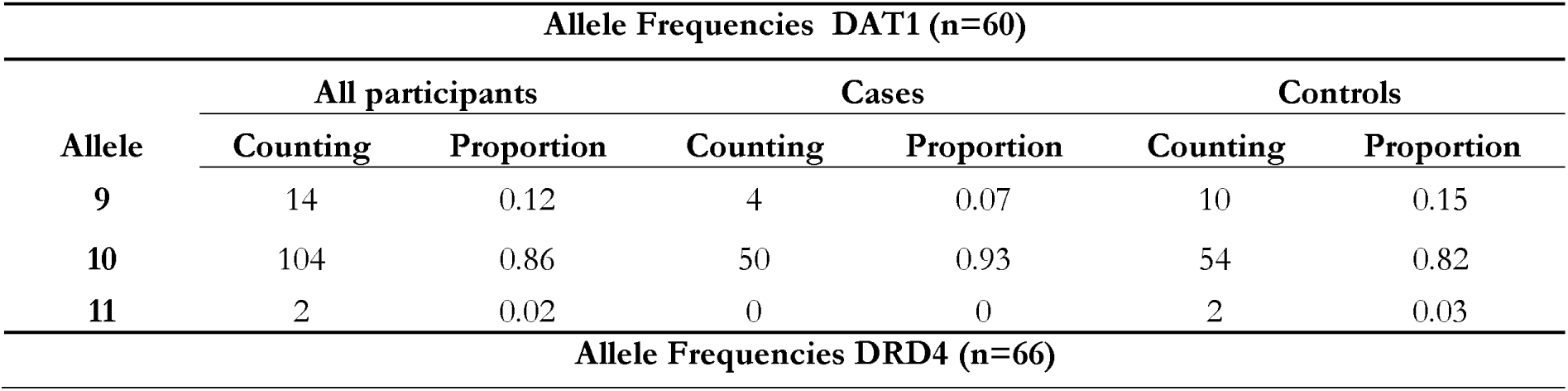
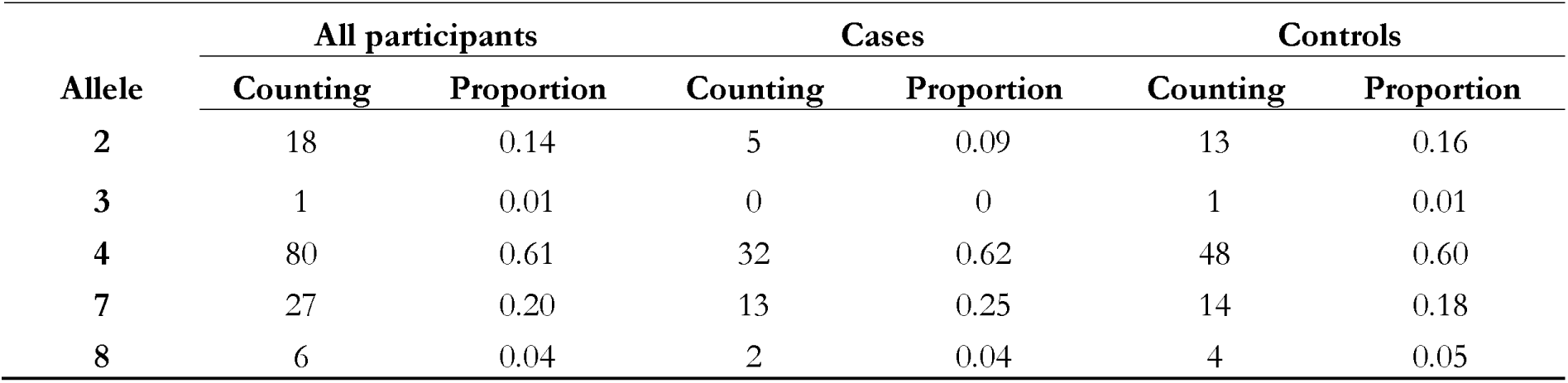
*Allele frequencies of polymorphic genes DAT1 DRD4*

As shown in Table 3, there is a higher frequency in the case group of 10-repeat allele (0.93) over the 9R allele, while 11R allele is not present. In contrast, the control group shows 10R allele as the highest (0.82), followed by allele 9 and, with a lower frequency, allele 11 (only 2 times). In the case of the VNTR exon III for DRD4 polymorphism, as identified in Table 2, the 4R allele shows a higher general presence, with a similar frequency in the total population and in the two groups (general 0.61; cases 0.62 and control 0.60), sharing its distribution mainly with the 7R allele (general 0,20; cases and control 0.25 0,18). A comparison analysis, where cases and controls were included, was conducted with the aim of analyzing the odds ratio of polymorphisms in candidate genes and develop the disorder.

**Table 4.** *OR for each allele* of *gen DAT1, DRD4 and manifestation of ADHD.*

| DAT1 |  |  |  | DRD4 |  |  |  |
| --- | --- | --- | --- | --- | --- | --- | --- |
| Allele | OR | IC (95%) | p-valor | Allele | OR | IC (95%) | p-valor |
| 9R | 0,45 | (0,132 – 1,518) | 0,19 | 2R | 0.548 | (0.183 , 1.642) | 0.278 |
| 10R | 2,78 | (0,841 - 9,178) | 0,08 | 4R | 1.067 | (0.521 , 2.182) | 0.86 |
| 11R | 0,59 | (0,052 – 6,715) | 0,57 | 7R | 1.571 | (0.67 , 3.685) | 0.297 |
|  |  |  |  | 8R | 0.76 | (0.134 , 4.31) | 0.557 |

The analysis of risk for independent alleles and the factor presence-absence of the disorder (group cases vs. control), for DAT1, allele 10R, is the one with the highest positive risk value, with a smaller p value (p: 0.08) compared to 9R and 11R alleles (p: 0.19 and p: 0.57), but none of the alleles shows statistically relevant differences between groups. In the case of the marker for gene DRD4, there is positive risk for alleles 4 and 7; without significance. According to the p-values found, is evident the independence between a given allele and manifestation of ADHD (P: <0,05).

**Table 5.**
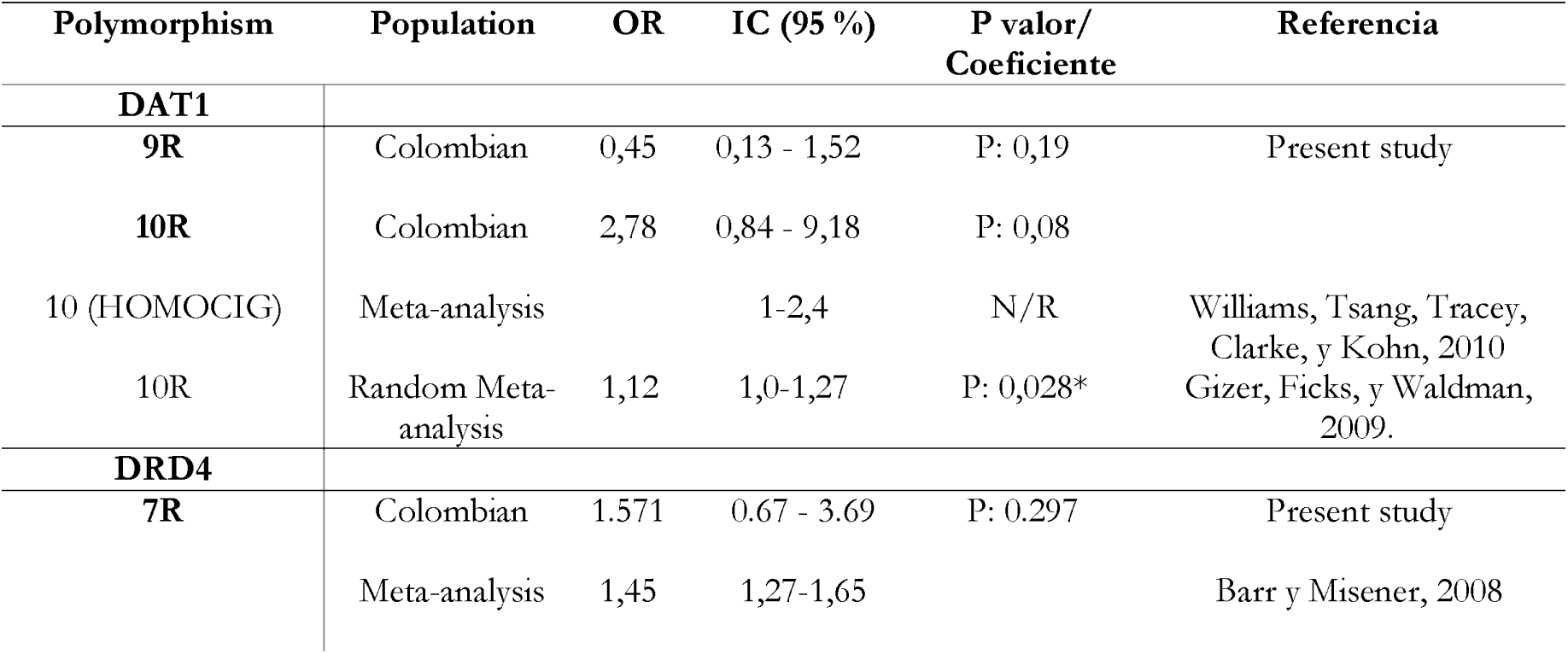
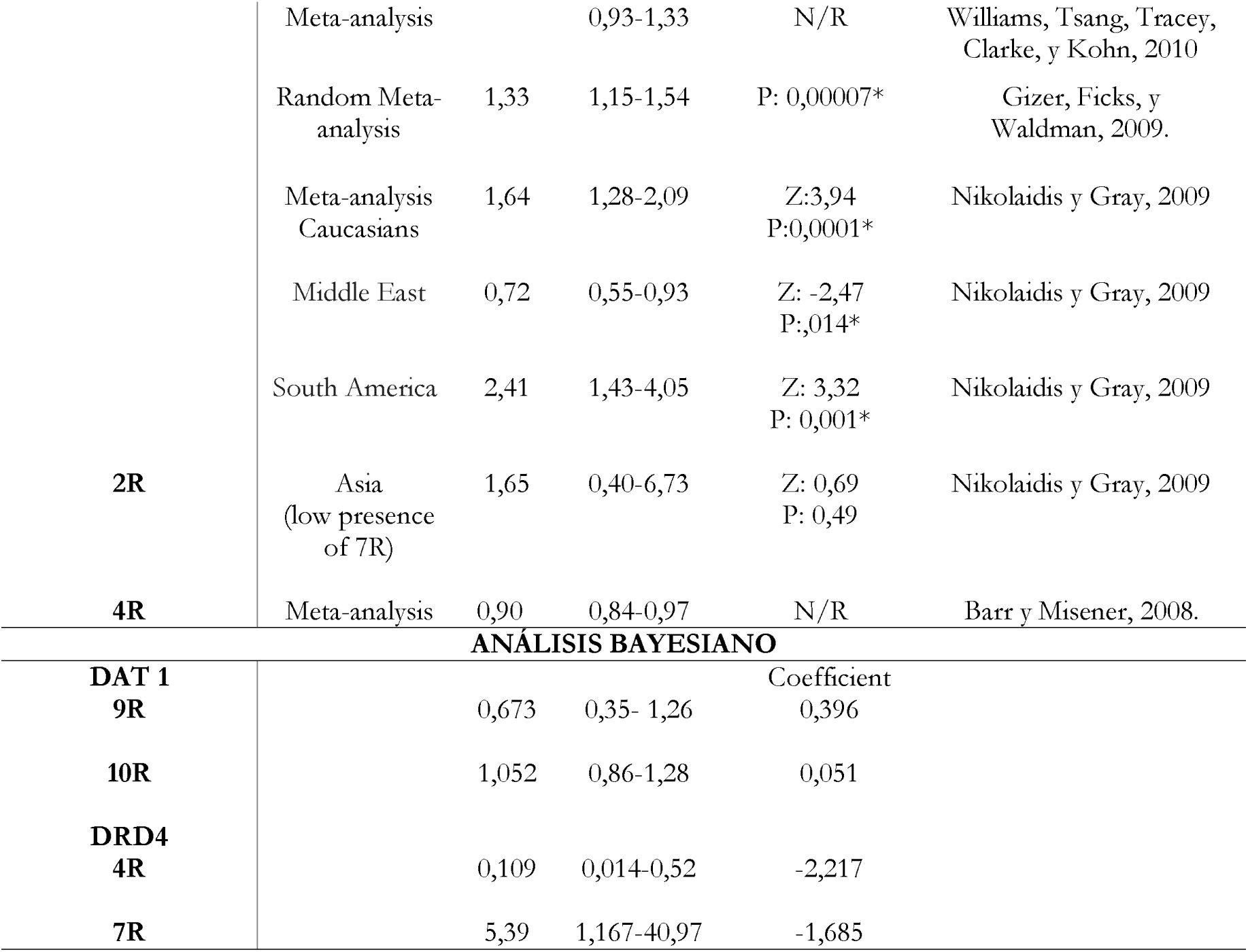
*Comparison of risk association studies and integrative Bayesian analysis model.*

When comparing the findings with other studies, consistency is evident in risk analysis of dopamine alleles in different populations, highlighting a significant risk in the South American population. The OR analysis shows no significant evidence between the groups, but with high risk, particularly for allele 10R in DAT1 and 7R in DRD4. Adjustment of a Bayesian logistic regression model was made to determine the influence of genetic markers on the probability of pathology with the estimates given in Table 5. Results shows that for gene DRD4 the risk associated with the presence of a 7R allele is five times higher, over those who do not have it.

For DAT1, despite a priori information considered for this study, there is no evidence to establish the existence of significant differences between the risk alleles, since high frequency of this variants in the population is an important element.

Considering the findings of likelihood of positive risk, an analysis to link more specifically alleles with cognitive functioning is proposed after identifying possible candidates for endophenotypes.

**Table 6.** *Analysis of the independence of cognitive variables from other genotypes vs. alleles for the marker of gene DAT1 and DRD4.*

| Pruebas Cognitivas | Presencia alelos frecuentes<br>Test Exacto de Fisher (1-sided)<br>p: <0,05 |  |  |  |
| --- | --- | --- | --- | --- |
|  | DAT 1 |  | DRD4 |  |
|  | 9R | 10R | 4R | 7R |
| Total IQ | 0,54 | 0,659 | 0,24 | 0,12 |
| Verbal Comprehension | 0,61 | 0,825 | 0,68 | 0,06 |
| Visual Spatial Index | 0,54 | 0,698 | 0,18 | 0,42 |
| Working Memory Index | 0,41 | 0,549 | 0,29 | 0,22 |
| Processing Speed Index | 0,58 | 0,33 | 0,14 | 0,03* |
| Coding | 0,46 | 0,38 | 0,10 | 0,21 |
| Symbol Search | 0,17 | 0,49 | 0,13 | 0,01* |
| Picture completion | 0,64 | 0,58 | 0,25 | 0,29 |
| Digit Span total | 0,36 | 0,69 | 0,29 | 0,53 |
| Letters Cancellation ENI | 0,29 | 0,58 | 0,37 | 0,33 |
| Pictures Cancellation ENI | 0,40 | 0,41 | 0,2 | 0,38 |
| Stroop interference score | 0,63 | 0,91 | 0,46 | 0,05 |

The DAT1 alleles does not show significant differences for the performance on cognitive tests. For the DRD4 markers, when considering the most frequent allele, no difference was evident in cognitive performance among those individuals who have the 4R allele and those with other alleles. Considering the 7R allele, significant differences are found in the performance for speed processing and symbol search tasks.

**Table 7.** *Association risk analysis (RR) for the presence of allele 7 and performance on cognitive tests.*

| Processing Speed Index WISC IV |  |  |  |
| --- | --- | --- | --- |
| Allele | RR | CI (95 %) |  |
|  |  | Minimum | Maximum |
| 7R | 2,302* | 1,070 | 4,952 |
| Symbol Search WISC IV |  |  |  |
| Alelo | RR | CI (95%) |  |
|  |  | Minimum | Maximum |
| 7R | 3,553* | 1,271 | 9,932 |

Based on the calculation of risk odds, in the presence of at least one 7R allele, it is evident that this marker is a risk factor for low performance on processing speed index and the search symbols task (considering the RR> 1 values), while having another allelic variant is a protective factor (OR: 0,607 IC: 0,388-0,952 p: 0,011).

## Discussion

By analyzing dopamine genes DAT1 and DRD4 in the study population, similar trends to those found in the general population are identified. In the case of 40-bp VNTR for the 3’UTR region in chromosome 5p15.3 (SLC6A3) of DAT1, several studies have shown that for the 10R allele, high frequencies are observed in the world’s population, and in the case of the Latin American population, this is close to ∼1.0, particularly in indigenous (Vieyra et al., 2003; Kang, Palmatier y Kidd, 1999; Carrasco et al, 2004) and Chilean population (0.81 and in cases and 0.70% in controls) (Carrasco et al., 2004), or in intermediate frequencies between European and indigenous population. However, a higher frequency is sustained in cases (ADHD), although without statistical differences between groups (Vieyra et al., 2003).

The high presence of this variant in the general population hinders the analysis of association between the polymorphism and attentional disorder (Kang, et al, 1999; Vieyra et al., 2003), even more so considering the frequency of occurrence of allele 10 in South American communities; nevertheless, this could be regarded as a factor that represents a characteristic of the population functioning and, in turn, a potential risk factor considering interaction with the environment.

The VNTR polymorphism in the dopamine receptor gene type 4 (DRD4) varies in relation to ethnicity and geographical distribution. In this research, a distribution, where the most frequent allele is 4W with 0.61 (cases and controls combined), which is similar to the values found globally with an allele frequency of 64.3%, appears in the population with frequencies in a range from 0.16 to 0.96 (Wang, Ding, Flodman, et al, 2004). The 7R allele, in the case of the Americas, appears more frequently (48.3%) than what has been identified (0.20%). Regarding the reports by Chang Kidd, Livak, Pakstis and Kidd (1996) about the Colombian population, 7R allele has a greater appearance of about 0.62, followed by alleles 4R at a rate of 0.23, which slightly differs from what is described here. However, the possibility of finding important events for the probability of risk associated with ADHD (OR: 1,571; CI at 95%: 0.67 - 3.69) is consistent with findings in different populations (Barr & Misener, 2008, Williams, Tsang, Tracey, Clarke, y Kohn, 2010, Gizer, Ficks, y Waldman, 2009, Nikolaidis y Gray, 2009), particularly, when the probability of risk is confirmed through a Bayesian analysis (OR: 5.39 IC 95%: 1.167 – 40.97, coefficient: −1.685), indicating a probability five times greater of presenting ADHD in the presence of such variant.

Generating explanations of phenomena such as ADHD requires an understanding of the complex dynamics between biological, cognitive and behavioral factors, and their relationship with the environment. In this sense, the search for possible relationships or interaction points when trying to determine the interaction between the presence of a polymorphism and the performance on cognitive tasks, as in this case, demonstrate the difference between children with ADHD and their peers without the diagnosis. Once the characteristics of cognitive performance are associated with genetic variants, we have that the presence of at least one allele of the 7R dopamine receptor (DRD4) implies a risk towards low performance in tasks linked to processing speed (VP index RR: 3.93 IC 95%: 1.114-13.851 p: 0.031) and symbol search (RR: 5.85 IC 95%: 1.464-23.377 p: 0.011).

The D4 receptor (DRD4) is primarily expressed in pyramidal neurons and interneurons in the prefrontal cortex, but there is also evidence of localization of DRD4 in medium spiny neurons in the basal ganglia (striatum and nucleus accumbens), and in the limbic system and the thalamus rodent. High densities in neuronal packages, known as striosomes, in striatal neurons on rodents (also called matrix) have been found. The estriosoma receives inputs from the cingulate cortex and thalamus, indicating an important role in the DRD4 for regulation of the cortico-striatal-thalamic loop (Currier, Grandy, Gerhardt y Glaser, 2009). Thus, DRD4 is expressed within the mesocorticolimbic dopamine pathway, which originates in the ventral tegmental area (VTA) and innervates the olfactory tubercle, the nucleus accumbens, the septum, the amygdala and the adjacent cortical structures, such as the entorhinal, perirhinal, and pyriform cortex, the medial prefrontal and, more importantly, the anterior cingulate (aMCC) (Taylor & Creese, 2002).

According to the model by Posner and Petersen (1990), the anterior attentional network/system or detection subsystem, consists of the anterior cingulate cortex (aMCC) and the dorsolateral prefrontal cortex, which play an important cognitive role involved in self-regulation, and that may respond to the detection of targets in addition to information being processed (Posner y Rothbart, 2009; Bush, 2010). The aMCCA has a role in modulating the activity in divided attention tasks and support the new processing (Bush, 2010). Similarly, it would be involved in maintaining voluntary alertness during tasks (Posner y Rothbart, 2009), interacting with impaired functioning in ADHD. These risk ratios establish that, those with allele 7R, have a higher risk of low performance (less than 1DT) in tests related to processing speed and selective attention within a time limit, which implies sustained attention.

A study based on reaction times in different cognitive tasks, carried out with 245 Caucasian individuals, found that, in a non-clinical population, participants with allele 7R showed slower responses, regardless of sex, which was not caused by fatigue (P = 0.0001). This occurred without association to a specific task, which relates to a deficit in the general attentional processing (Szekely et al., 2010).

It was also estimated that the DRD4 7R allele is more associated with inattention than with hyperactivity—possibly DAT1 would be more linked to this characteristic (Diamond, 2005)—, as well as with an increase in commission errors in individuals with ADHD when executing tasks of sustained attention (Greene, Braet, Johnson y Bellgrove, 2008; Sheppard & Vernon, 2008; Szekely et al., 2010). This allele has also been associated with a style of impulsive and inexact responses in ADHD, regardless of the severity of the symptoms (Greene, Braet, Johnson y Bellgrove, 2008), a factor that could be related to the risk of the Stroop Test, where having an allele different to 7R is a protective factor (RR: 0.842 IC: 0.69-1.02 p: 0.53), without reaching a level of significance of 5%, but of 10%. By comparing these findings with the performance of children with ADHD, it is observed that the flaws would be directed towards problems in the voluntary control of attention at the time to distinguish the particular characteristics of the relevant stimulus, those that become distractors (signal/noise) (Williams, Tsang, Clarke y Kohn, 2010; Posner y Rothbart, 2009; Wood et al., 2011). This could be linked directly with low alertness affecting the voluntary control system (self-regulation) all the time, which would be reflected in a general decline in processing speed, attentional lapses and low performance over time (Williams et al., 2010; Bellgrove, O’Connell and Vance, 2008; Szekely et al., 2010).

When considering the presence of the receptor DRD4 within regions related to previous attentional or executive network (Posner y Rothbart, 2009, Bush, 2010), particularly in pyramidal neurons within aMCC, we can find a possibility of allele 7R slowing down dopaminergic modulation exerted on this system, directly involved in voluntary control of alertness and attentional performance. *In vivo* studies with animal models, using wild type mice (DRD4 +/+), heterozygotes (+/− DRD4) and knockout (KO) (DRD4 − / −), Rubinstein and colleagues (2001) presented evidence through immunohistochemical, electrophysiological, pharmacological and ultrastructural methods, in which the KO DRD4 −/− mouse exhibited cortical hyperexcitability. These results were consistent with the fact that the activation of the DRD4 has an inhibitory effect on pyramidal neurons containing glutamate in the frontal cortex. Therefore, in cases where there is no presence of the DRD4 receptor, as in the KO model, there is an increase in the firing of neurons containing glutamate and a significant slowing in the elimination of this neurotransmitter in the striatum, indicating a possible role for DRD4 in glutamate regulation in the striatal circuit, which implies changes in glutamatergic neurotransmission producing some sort of hyper-glutamatergic state (Currier et al., 2009).).

Lichter *et al.* (1993) propose a variation in receptor protein due to the change in the number of repetitions, manifested in three possible roles: 1. different cytoplasmic loops that change the conformation of the transmembrane domains and alter ligand binding; 2. the variation in the number of repetitions affects the translation of the signal, altering the interaction with G proteins downstream or other intracellular proteins; or 3. no functional consequences (Chang et al., 1996). Then, if we consider the effect described in the animal model with KO (Currier et al., 2009) mice, it could be said that the variant 7R in the DRD4 produces hypoactivity in the receptor, which means that there is no inhibitory effect on pyramidal glutamatergic cortical neurons, particularly those distributed in the aMCC, leading to cognitive manifestations described in participants diagnosed with ADHD on tasks related to processing speed, the effect of a failure in tasks related to anterior/executive attentional network (Posner y Rothbart, 2009, Bush, 2010), which is summarized in Figure 1.

**Figure 1.** *Explanatory model based on the risk association between allele DRD4 of the 7R gene and cognitive performance. A. Anatomical representation of the dopaminergic regulation system and the influence of the 7R allele on the anterior midcingulate cortex (aMCC). B. Hypothesis of correlation between anatomical and physiological processes associated with the functioning of the receptor from the VNTR with 7R allele and cognitive performance identified in ADHD.*

Dopaminergic modulation can increase the neuronal signal-noise ratio by boosting the signal to cushion the surrounding noise. It also displays an inverted U type influence, so that neural transmission can be optimized within a range, but can adversely affect performance at low or high levels, as in the case of a modulation decreased by allele 7R (Bush, 2010).

Nonetheless, it is important to consider the gene-environment interaction in this possible relation because, according to various descriptions, the environment can have a large influence on those with allele 7R by, for example, showing that those with this variant respond better to an intervention aimed at the importance of the parental role (Posner y Rothbart, 2009).

From this perspective, cognitive mediations aligned to the management of attentional, self-control from processing speed would imply that, as processing speed improves, memory span also improves. The speed implies that stimulus should not last much in the mind, reducing demand for working memory. This leads to co-variation of executive function and speed because both demonstrate more efficient neural processing and improved signal/noise rates. The search for a better function of the prefrontal cortex would facilitate the improvement of the signal/noise ratio for different neural regions, allowing speed and greater efficiency in cognitive processing (Diamond, 2005).

This proposal, then, becomes a possibility of building an explanatory bridge that connects different areas of knowledge, which may be basic (when including genetic variations) and integrative of different cognitive and neural functioning models to propose explanatory models that can really open up a whole new field applied toward the development of new etiological and interventional possibilities, based on the analysis of different levels of interaction between systems, which summarize the complexity of conditions such as Attention Deficit Disorder and Hyperactivity.

Clearly, further studies that include an increased sample size, greater control of variables related to comorbidity and severity of symptoms, are required, as well as using more specific cognitive measures (*e.g.* SSRT, direct response speed measures, reaction time, among others) in search of key elements for diagnosis and treatment, where the concept of endophenotype appears, becoming a valuable aspect for providing measurable markers close to the causes of ADHD (Graham, Seth & Coghill, 2007).

